# Formation of a swelling gel underlies a morphological transition in *Bacillus subtilis* biofilms

**DOI:** 10.64898/2026.02.20.707077

**Authors:** Ayantika Saha, Joshua M. Jones, Abigail Plummer, Joseph W. Larkin

**Affiliations:** Department of Physics, Boston University, Commonwealth Ave., Boston, 02215, MA, USA; Department of Mechanical Engineering, Boston University, Commonwealth Ave., Boston, 02215, MA, USA; Department of Biology, Boston University, Commonwealth Ave., Boston, 02215, MA, USA

**Keywords:** gelation, phase transition, microbial communities, living material, hydrogel swelling

## Abstract

Microbes across species and environments form biofilms, living materials composed of cells and extracellular polymers. Biofilm-dwelling cells benefit from emergent soft matter physics that sculpts three-dimensional morphologies and osmotically absorbs nutrients. Although biofilms are modeled as viscoelastic gels, the physical origins of the phase transition underlying their conversion from groups of cells to living gels have not been systematically investigated. Here, we show that *Bacillus subtilis* biofilms use polymer composition to tune their physical properties and drive gel formation. Using imaging and water immersion experiments with matrix knockout strains, we demonstrate the complementary roles of two polymers in this developmental transition: hydrophilic poly-***γ***-glutamate swells colonies by absorbing water and exopolysaccharides serve as effective cross-linkers, causing a sol-gel-like phase transition that imparts structural integrity. With matrix knockout co-culture biofilms, we independently modulate the production of each polymer and reveal a phase space of biofilm morphologies. Colonies that produce both polymers develop macroscopic wrinkles. A thin-film model predicts biofilm wrinkling from swelling-generated internal strain coupled to elasticity. The model reproduces the shape of our observed morphological phase diagram. Our results demonstrate that bacteria leverage gelation to vary their material properties and morphologies, with implications for microbial ecology and engineering living matter.

---

In many environments, microbial cells live in a self-produced polymeric matrix as biofilms[1]. Why has evolution so consistently prioritized this metabolically costly community structure? The soft matter physics of these communities gives cells new opportunities unavailable to solitary microbes, such as osmotically driven expansion [2, 3], enhanced nutrient capture[4], water retention[5], and formation of 3D wrinkled structures[6–9] that improve oxygen availability[10]. For biofilms to take advantage of these emergent physics, they must turn themselves into viscoelastic materials by secreting extracellular matrix (ECM) polymers, demonstrating a fundamental connection between polymeric phase transitions and multicellular communities. To understand the impact of polymeric phase transitions on microbial ecology and to exploit their unique properties, we must investigate their origins in ECM production and their consequences for biofilm properties such as morphology and environmental response. Characterizing and understanding this self-regulated phase transition will provide insight into the complexities of biofilm formation and function[5, 11], relevant to a wide range of industrial and medical applications[12, 13] as well as efforts to engineer living materials[14].

Previous studies have investigated how material properties impact biofilm growth and physiology[2, 7, 15–18], but they have generally not examined how colonies transition between distinct material phases depending on the abundance and physicochemical properties of ECM polymers, nor how those transitions regulate colony-level phenomena such as three-dimensional morphology and responses to perturbations. The soil bacterium *Bacillus subtilis* is an ideal system in which to investigate these questions. The species produces polymers with distinct physical properties, including exopolysaccharides (EPS, distinct from EPS as a general term for biofilm extracellular polymeric substances)[19], poly-*γ*-glutamic acid (PGA)[20], and several others[21]. PGA and EPS in particular have been shown to play key roles in biofilm material properties. PGA, being highly hydrophilic[22], absorbs water into biofilms, potentially producing locally fluid regions[15]. EPS production increases biofilm elasticity[6, 17] and recent results show that purified *B. subtilis* EPS exhibits a gel phase transition with increasing EPS concentration[19]. These results suggest that by synthesizing both PGA and EPS, biofilms could form swelling gels with distinct properties depending on the amount of each polymer synthesized.

Here we investigate phase behavior in *B. subtilis* biofilms, varying ECM production to tune the colony from a fluid droplet-like state to a cross-linked swelling gel that can form complex 3D morphologies. This biofilm phase space has not been previously explored due to difficulty in experimentally modulating ECM production and the use of *B. subtilis* strains that are deficient in PGA production—the commonly studied biofilm-forming *B. subtilis* strain NCIB3610 (3610 hereafter) produces little to no PGA under standard lab biofilm conditions[15], unlike many soil isolate strains[23]. However, we have recently found that a mutation in the plasmid-borne gene *rapP* inhibits PGA production in 3610. Deleting *rapP* or curing 3610 of *rapP* ‘s plasmid vector pBS32[24] restores the effect of PGA on *B. subtilis* biofilm formation[25]. Building on this finding, we create matrix knockouts in this PGA-producing strain and perform co-culture experiments inoculated at different colony-forming unit (CFU) ratios. By systematically modulating the proportion of PGA- and EPS-producing cells in biofilms, we navigate a wide range of polymer compositions and uncover a phase space of diverse biofilm morphologies and material properties.

We find that EPS production drives a gelation-like transition from colonies that completely dissolve in water to elastic films that remain intact, while PGA absorbs water into colonies and increases vertical thickness. We show that both PGA and EPS are necessary for the formation of macroscopic wrinkles. To model the effect of polymer production on biofilm morphology, we develop an elastic bilayer model that predicts colony wrinkling from the interplay of PGA-driven swelling and EPS-driven elasticity. By imaging co-culture colonies, we measure biofilm morphology across the PGA-EPS phase space and find a wrinkling transition that qualitatively matches the shape predicted by our model: only colonies that produce both polymers form large wrinkles. Our results show that soil-dwelling bacteria take advantage of a gelation transition to form fluid-absorbing, living materials.

## Self-secreted polymers PGA and EPS shape biofilm surface morphology

The surface topography of bacterial biofilms is heavily influenced by the composition of the ECM. Though PGA and EPS have been shown to qualitatively alter the physics of *B. subtilis* colonies[15, 19], their coupled impact on biofilm morphology is poorly understood. To investigate these relationships, we created four strains that each either produce or do not produce PGA and EPS (see Methods). Our wild-type (WT) *B. subtilis* strain (NCIB3610 pBS32^0^)[24] synthesizes both PGA and EPS (PGA^+^, EPS^+^). Δ*pgsB* (NCIB3610 pBS32^0^ Δ*pgsB*) produces EPS but not PGA (PGA^−^, EPS^+^); Δ*eps* (NCIB3610 pBS32^0^ Δ*epsA* − *O*) produces PGA but not EPS (PGA^+^, EPS^−^), and the double knockout Δ*pgsB* Δ*eps* (NCIB3610 pBS32^0^ Δ*epsA* − *O*) produces neither polymer (PGA^−^, EPS^−^). We refer to these as the four “base strains”.

Following 48 hours of incubation on biofilm-stimulating MSgg media[26] (see Methods), each strain formed morphologically distinct colonies. We visualized lateral morphology with stereomicroscopy (Fig. 1a-d, see Fig. S1a-d for images at 24 hours; see Methods) and cross-sectional morphology using optical coherence tomography (OCT; see Methods) [27] (Fig. 1e-h). Biofilms deficient in both PGA and EPS (Δ*pgsB* Δ*eps*) formed thin, smooth colonies (Fig. 1a, e). Colonies synthesizing EPS but not PGA (Δ*pgsB*) exhibited a modest increase in thickness and developed microscale surface ridges (Fig. 1b, f). Biofilms producing only PGA but no EPS (Δ*eps*) developed a markedly thicker morphology (*>* 100 *µ*m), with a smooth, droplet-like profile that glistened (Fig. 1c, g). WT biofilms, which produce both PGA and EPS, formed thick colonies characterized by a complex surface morphology with 100 *µ*m-scale wrinkles (Fig. 1d, h, see Fig. S4a, b, e, f for measurements on the common biofilm strain 3610 and PS-216, a biofilm-forming isolate lacking the pBS32 plasmid[28]). To summarize, biofilms that produced PGA were notably thicker than those that did not, but only biofilms that produced both PGA and EPS formed dramatic, macroscopic wrinkles. To investigate how these ECM polymers alter biofilm material properties and enable wrinkling, we imaged the four base strains early in biofilm development as EPS and PGA were produced.

**Fig. 1.**
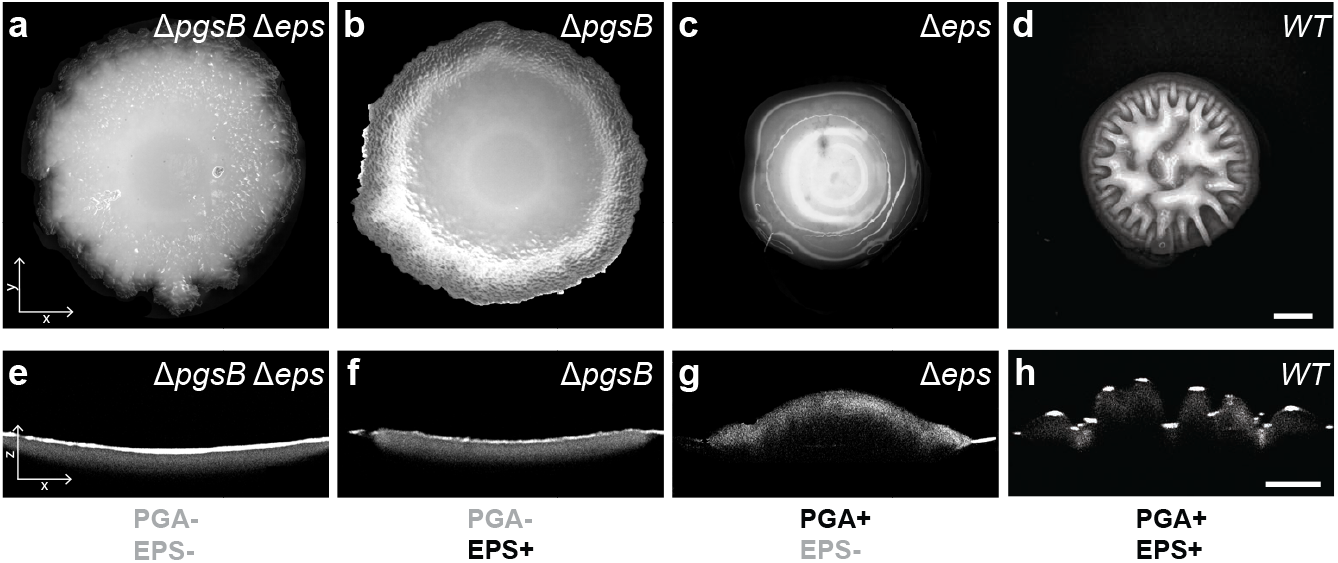
Polymer composition shapes biofilm surface morphology. **a–d**, Top-down images of 48-hour-old *B. subtilis* biofilms formed by strains with differential production of PGA and EPS polymers (polymer production in figure bottom row): **a**, Δ*pgsB* Δ*eps* (neither polymer), **b**, Δ*pgsB* (EPS only), **c**, Δ*eps* (PGA only), **d**, WT (both polymers). **e-h**, Corresponding *xz* cross-sectional profiles measured by optical coherence tomography. Edge curvature is from the agarose substrate. Scale bar in both rows, 500 *µ*m.

## PGA is required for fluid micro-environments in developing biofilms

We hypothesized that the swollen morphology observed in PGA-producing biofilms (Fig. 1c, d, g, h) arises from PGA-mediated water absorption from the underlying agarose substrate, and that PGA-producing biofilms would form fluid or fluid-like regions[15]. To test this hypothesis, we acquired high-speed phase-contrast movies of developing colonies of the four polymer base strains (Fig. 2a-h), in which we could observe whether the biofilms exhibited rapid internal motion consistent with motile bacteria navigating a fluid environment[29]. Initially, all four strains looked identical (Fig. S2e-h). After approximately 10 hours, coinciding with initial expression of PGA ligase *pgsB* (Fig. S2a), the WT biofilm exhibited a distinct phase-dark annular region at the colony periphery (Fig. 2d), while the Δ*eps* strain displayed a similarly darkened region throughout the colony interior (Fig. 2c). In contrast, both PGA-deficient strains (Δ*pgsB* and Δ*pgsB* Δ*eps*) lacked such phase-dark features (Fig. 2a, b). We hypothesized that the phase-dark regions of PGA-producing biofilms were fluid micro-environments based on previous observations of 3610 biofilms grown under PGA-promoting conditions[15]. To determine if these regions were fluid, we performed high-frame-rate imaging in the areas of interest (yellow dotted boxes in Fig. 2a-d) and tracked feature motion over short time scales (∼60 seconds) (see Methods). Due to the resolution limitations of 10X phase-contrast microscopy, it was not possible to discern whether the observed features were individual cells or cellular clusters. We visualized feature trajectories using color-coded speed mapping (Fig. 2e-h; Supplementary Videos V1-V4) at *t* = 0 and 10 hours. We expected that fluid regions would support rapid motion due to active cell motility or passive diffusion, whereas solid regions would exhibit little to no detectable motion in high frame-rate videos. As controls, we tracked feature motion in a droplet of fresh motile cell culture before (wet) and after (dry) evaporation. Consistent with our expectations, we observed high-speed motion in the wet droplet and negligible motion in the dried droplet (Supplementary Videos V5, V6, and Fig. S2b-d for positive and negative controls).

**Fig. 2.**
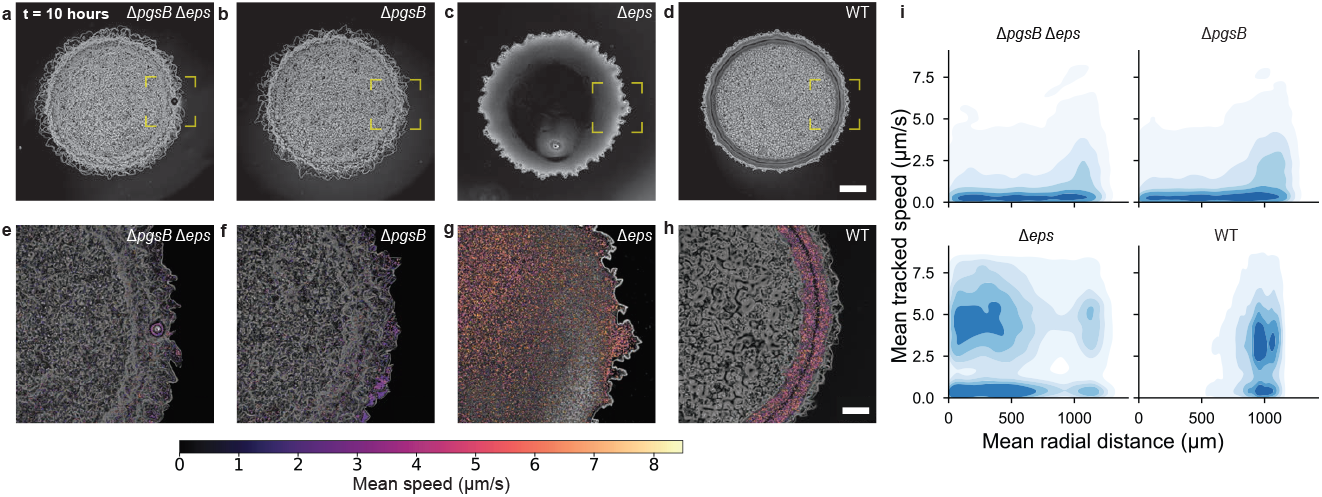
High-speed cellular motion tracking reveals fluid regions within PGA-producing biofilms. **a–d**, Phase-contrast images of **a**, Δ*pgsB* Δ*eps*, **b**, Δ*pgsB*, **c**, Δ*eps*, and **d**, WT after 10 hours show phase-dark areas in PGA-producing biofilms, scale bar 500 *µ*m. **e-h**, Speed-coded particle trajectories (*N* ≈ 17000 particles) from 60-second phase movies show sustained motility in WT and Δ*eps* but not in PGA-deficient Δ*pgsB* and Δ*pgsB* Δ*eps*. Scale bar 200 *µ*m. **i**, Kernel density estimates (KDEs) of mean tracked cell speeds and radial trajectory distributions confirm the presence of fast, directed cell motion in strains producing PGA. WT, which produces PGA and EPS, exhibits cell motility only at the colony edge.

At *t* = 0 hours, the phase time-lapse video yielded identical speed and trajectory distributions across strains (see Fig. S2e-i). All four base strains exhibited little to no detectable motion. By *t* = 10 hours, however, the strains were distinct. We observed rapid feature motion in PGA-producing biofilms (WT and Δ*eps*, Fig. 2g, h, i bottom). and little to no motion in PGA non-producers (Δ*pgsB* and Δ*pgsB* Δ*eps*, Fig 2e, f, i top). In PGA-producing strains, the spatial distribution of tracked particle trajectories varied depending on whether the strain produced EPS. In WT biofilms (EPS^+^), rapid motion was confined to the outer annulus (Fig. 2h, i bottom right), while in Δ*eps*, it occurred throughout the colony (Fig. 2g, i bottom left). These data are consistent with our initial hypotheses that PGA absorbs water from the agarose substrate, while EPS triggers biofilm solidification. PGA is necessary to generate fluid regions, but colonies that produce both PGA and EPS only exhibit fluidity at their outer edges. To further test these hypotheses, we investigated the physical effect of each polymer in isolation.

## Biofilm height scales with the fraction of PGA producers in the biofilm

To isolate the effect of PGA and EPS on biofilm morphology, we performed experiments with co-culture biofilms of the four base strains (see Methods). By inoculating colonies with different ratios of each strain, we can independently vary colony polymer production and observe the effect on biofilm physics and morphology. We first examined the effect of modulating PGA production in the absence of EPS by growing co-culture biofilms of PGA-producing Δ*eps* (PGA^+^, EPS^−^); and non-PGA-producing Δ*pgsB* Δ*eps* (PGA^−^, EPS^−^). Matrix production may incur a fitness cost [30], which could affect the growth rates of different strains in co-cultures [31]. We therefore needed to verify that the co-culture ratios remain relatively stable over the duration of experiments in order to correlate initial CFU ratios with observed biofilm properties. We measured the CFUs of each strain with an antibiotic-based plating assay (see Methods) and found that co-culture biofilms approximately maintained their initial PGA^+^ CFU ratios over 48 hours (Fig. 3c). We consequently assume that PGA production increases monotonically with the fraction of initial PGA^+^ CFUs.

**Fig. 3.**
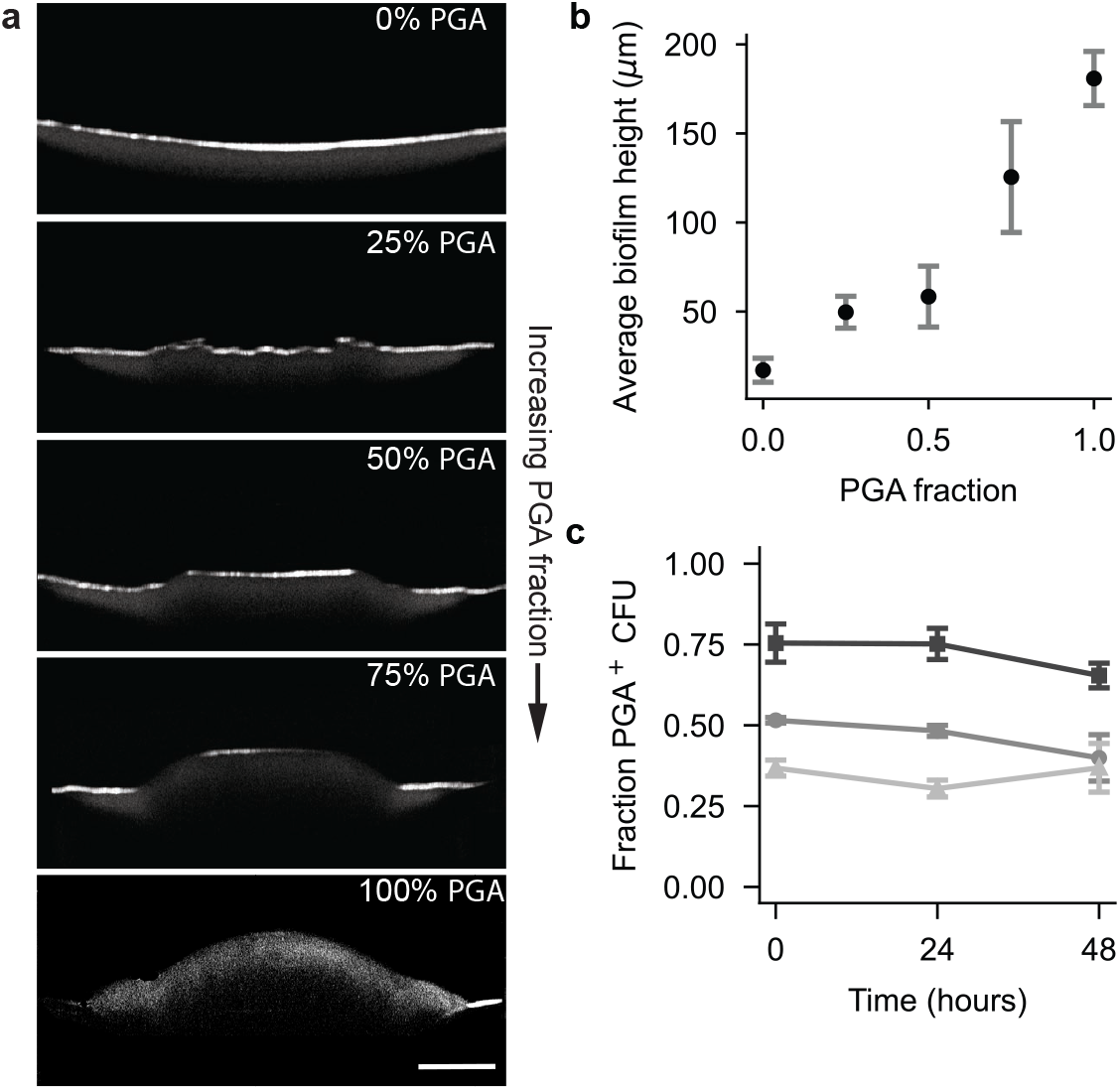
PGA-induced biofilm swelling determines biofilm thickness. **a**, *xz*-OCT cross-sections of co-culture biofilms with increasing fractions of PGA–producing cells. Scale bar, 500 *µ*m. **b**, Maximum biofilm height increases monotonically with PGA producer fraction, indicating fluid uptake–driven volumetric expansion. **c**, Proportion of Δ*eps* (i.e. PGA-producing) CFUs over time in co-culture biofilms with three different initial genotypic ratios of Δ*eps* and Δ*pgsB* Δ*eps* cells, 0.75:0.25 (squares), 0.5:0.5 (circles), and 0.25:0.75 (triangles). In b and c, points represent average ± S.D. from three independent replicates (*n* = 3)

In OCT experiments, we observed that PGA-producing biofilms are substantially thicker than non-producers (Fig. 1e-h). This observation, coupled with PGA production creating fluid environments in biofilms (Fig. 2), led us to hypothesize that an increase in PGA production leads to an increase in fluid absorption, which swells biofilms. The resulting increase in biofilm volume is then expected to increase biofilm height. Increases in the biofilm footprint depend additionally on the biofilm-substrate adhesion, which is more nuanced. To test this hypothesis, we acquired OCT cross-sections of co-culture biofilms after 48 hours of growth (Fig. 3a). We found that average biofilm height increased monotonically with the initial PGA producer fraction (Fig. 3b), consistent with our hypothesis of PGA-induced fluid absorption. We next examined the effect of changing EPS production on biofilm material properties.

## EPS drives gel formation in biofilms

We observed that EPS production inhibited the complete fluidization of biofilms (Fig. 2d, h). This led us to ask what physical mechanism could underlie this observation. Given that purified EPS is known to form gel-like polymer networks[19], we hypothesized that EPS-producing biofilms resist fluidization by forming a cross-linked gel where EPS acts as a structural scaffold. To test this hypothesis, we used a common assay for gel formation[32]: we immersed biofilms in water, the primary solvent for the biofilm polymer network. In this experiment, a microscopically cross-linked biofilm should retain structural integrity upon solvent addition, whereas a non-cross-linked, fluid-like biofilm should dissolve. Consistent with our hypothesis, both EPS-producing strains–WT (PGA^+^, EPS^+^) and Δ*pgsB* (PGA^−^, EPS^+^)–remained structurally intact upon water immersion for 30 minutes. In contrast, biofilms of the non-EPS-producing mutants–Δ*eps* (PGA^+^, EPS^−^) and Δ*pgsB* Δ*eps* (PGA^−^, EPS^−^)–dissolved in water (Fig. 4a). Moreover, we observed that WT biofilms swelled substantially in water, while Δ*pgsB* biofilms remained structurally intact but did not increase in volume (Fig. 4a, b, see Fig. S4c,d for the same measurements on 3610 and PS-216). To quantify the response of each strain to immersion, we measured the adhered biofilm area and height before (*A*_*i*_, *h*_*i*_) and after (*A*_*f*_, *h*_*f*_) water addition (see Methods). We then computed the fold change in each geometric quantity (*A*_*f*_ */A*_*i*_ and *h*_*f*_ */h*_*i*_). *A*_*f*_ */A*_*i*_ ≈ 0 indicates that biofilms dissolve in water and *A*_*f*_ */A*_*i*_ ≈ 1 indicates that they remain intact, as the area of adhered, residual biomass is roughly the same before and after water addition. *h*_*f*_ */h*_*i*_ *>* 1 indicates that a biofilm swells in water. The results for the four base strains are plotted in Fig. 4c. WT swelled, Δ*pgsB* remained intact but slightly shriveled, and both non-EPS strains dissolved entirely. These findings support the hypothesis that EPS polymers facilitate the formation of a cross-linked matrix with structural integrity that prevents biofilm dissolution in water. Additionally, the swelling of WT biofilms upon hydration is consistent with the swelling behavior of cross-linked hydrogels [33], and thus reinforces our earlier conclusion that PGA acts as a hygroscopic agent, promoting water absorption into the biofilm matrix.

**Fig. 4.**
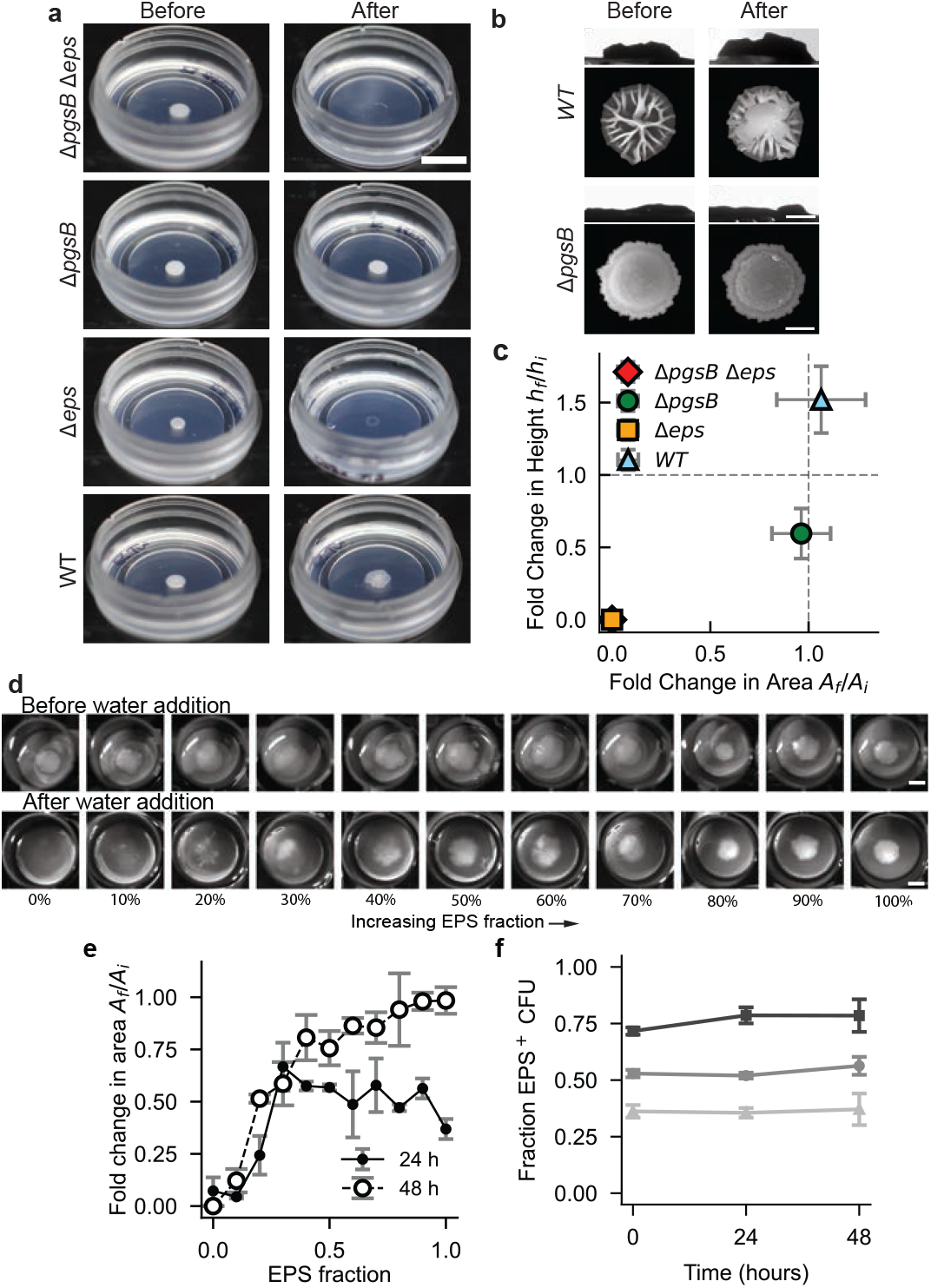
EPS production drives gelation. **a**, Representative images showing the state of biofilms before and after immersion in water. Biofilms containing EPS remain intact (Δ*pgsB*) or swell (*WT*), while those lacking EPS (Δ*eps* and Δ*pgsB* Δ*eps*) dissolve completely. Scale bar 5 mm. **b**, Top-down stereoscope images (top) and side-view photos (bottom) showing WT and Δ*pgsB*) of biofilms before and after water immersion demonstrate PGA-induced swelling in WT and no apparent height change in Δ*pgsB* . Scale bars 1 mm for top-down images and 2 mm for side-view images. **c**, Quantification of water immersion response (fold change in top-down area on *x*-axis and height on y-axis) for the four base strains shows complete dissolution for non-EPS producing strains, no significant change in area for Δ*pgsB*, and swelling in WT. **d**, Stereoscopic images before (top) and after (bottom) water immersion for 48-hour co-culture biofilms (Δ*pgsB* and Δ*pgspB* Δ*eps*) of varying EPS production show transition from dissolution to remaining intact. Scale bar (top and bottom) 5 mm. **e**, Ratio of retained area to actual biofilm area after DI H_2_O addition of 24-hours-old and 48-hours-old biofilms. **f**, Proportion of Δ*pgsB* (i.e. EPS-producing) CFUs over time in co-culture biofilms with three different initial genotypic ratios of Δ*pgsB* and Δ*pgsB* Δ*eps* cells, 0.75:0.25 (squares), 0.5:0.5 (circles), and 0.25:0.75 (triangles). In c, e, and f, points represent average ± S.D. from three independent replicates (*n* = 3).

Many gel formation processes can be described as a percolation phase transition close to the gel point [32]. If EPS-induced gelation occurs in biofilms, we would anticipate a sharp transition between biofilms dissolving to biofilms remaining intact as a function of EPS concentration. To investigate this hypothesis, we prepared co-culture biofilms with different initial proportions of Δ*pgsB* (PGA^−^, EPS^+^) and Δ*pgsB* Δ*eps* (PGA^−^, EPS^−^) to vary EPS producer fraction in the absence of PGA polymers. We validated that the relative abundance of Δ*pgsB* cells closely matched the initial inoculation ratio over 48 hours (Fig. 4f). As we increased the proportion of EPS^+^ CFUs, biofilms sharply transitioned from complete or near-complete dissolution in water to remaining intact, as shown in the series of before-and-after stereoscope images in Fig. 4d. We quantified dissolution by computing *A*_*f*_ */A*_*i*_ for both 24-hour-old and 48-hour-old biofilms (Fig. 4e). We found that the dissolution-resistant area exhibited a step-function-like transition consistent with a gelation phase transition. Beyond a critical initial proportion of EPS^+^ cells, biofilms formed water-stable networks across whole colonies. At low EPS^+^ proportion, there is insufficient cross-linking to connect the colonies macroscopically, resulting in dissolution. For the 48-hour-old biofilms, we observed a shift of the transition point to a lower critical EPS producer fraction, as well as a higher *A*_*f*_ */A*_*i*_ ratio. This may occur due to the accumulation of EPS polymers as EPS^+^ cells continue to synthesize EPS between 24 and 48 hours, which correspondingly lowers the critical CFU threshold for preventing dissolution.

## Combined modulation of PGA and EPS reveals a morphological transition in biofilm architecture

Our experiments suggest that PGA swells the biofilm while EPS confers stiffness via a sol-gel phase transition (Fig. 5a). To assess whether these effects can account for the biofilm morphologies we observed in the base strains (Fig. 1), we adapted a standard wrinkling threshold calculation for a compressed thin film adhered to a thick compliant substrate to predict biofilm wrinkling based on the fraction of PGA-producing and EPS-producing CFUs in a colony (*f*_*PGA*_ and *f*_*EPS*_, respectively, see Supplementary Information for model details). As the fraction of PGA-producing CFUs in a colony (*f*_*PGA*_) is increased, the biofilm height is observed to increase monotonically (Fig. 3b). Moreover, the in-plane contact radius is found to decrease with increasing fraction of PGA-producing CFUs, possibly due to increased adhesion between the biofilm and the substrate. This height increment, coupled with a decrease in the base radius implies the presence of in-plane compressive stress because the stress-free state of a swollen biofilm has a greater in-plane area [7, 34–36]. On the other hand, as the fraction of the EPS-producing CFUs (*f*_*EPS*_) is increased, the crosslinking, and presumably the stiffness *E*_*f*_, of the biofilm increases. The standard elastic bilayer calculation for the onset of wrinkling requires that both the film and the substrate are elastic, the film is thin and much stiffer than the substrate, and that the film experiences in-plane compression relative to the substrate [37–41]. If these conditions are met, wrinkling occurs when the mechanical strain of the film measured relative to its stress-free state exceeds a threshold value that scales as a function of the stiffness of the colony. PGA or EPS could therefore cause wrinkling by either increasing compressive strain or by decreasing the wrinkling threshold, respectively. The model predicts a phase boundary that separates the morphological phase space into two regions depending on *f*_*PGA*_ and *f*_*EPS*_ - one where smooth, flat morphology is favored by the elastic energy and one where a wrinkled morphology is preferred (Fig. 5b). To test whether the observed morphology of *B. subtilis* biofilms is consistent with this model as the PGA and EPS fractions are varied, we again turned to co-culture experiments.

**Fig. 5.**
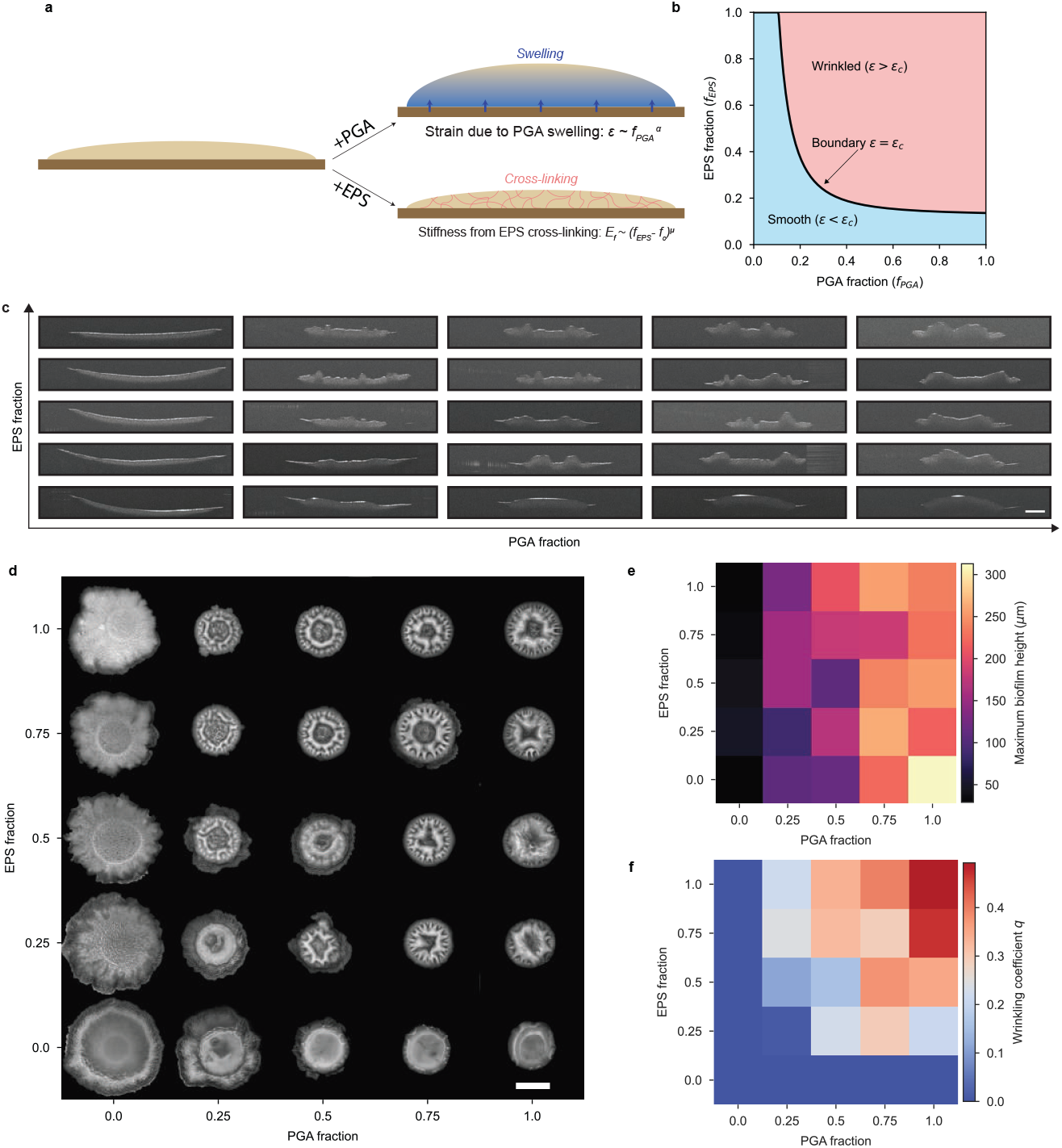
Bilayer thin film model predicts wrinkling transition via mechanical interplay between PGA–induced swelling and EPS-mediated gelation. **a**, Schematic summarizing the mechanical interplay between strain induced by PGA–driven swelling and EPS-mediated gelation providing elasticity. The in-plane compression in an elastic thin bilayer results in wrinkled surface morphology. **b**, Simulated phase diagram of biofilm morphology obtained by setting *α* = 2, *µ* = 1.5, *K* = 0.01 and *f*_*c*_ = 0.126 shows smooth and wrinkled phases as a function of PGA and EPS production. **c**, OCT-derived *xz*-profiles of biofilms with increasing fractions of PGA and EPS–producing cells (0–100%). Scale bar 1 mm. **d**, Top-down images of co-culture biofilms across the PGA-EPS phase space with initial PGA-producing CFU proportion increasingly along the *x*-axis and initial EPS-producing CFU proportion increasing along the *y*-axis. Scale bar 2 mm. **e**, Heatmap of maxi\mum heights quantified from segmented surfaces in (c), with PGA and EPS fractions mapped along the *x* and *y* axes, respectively. **f**, Heatmap of wrinkling coefficients quantified from segmented images in (d), with PGA and EPS fractions mapped along the *x* and *y* axes, respectively.

We varied both PGA and EPS in biofilms with co-culture colonies of all four base strains. By inoculating CFUs of the strains at different ratios, we could independently modulate the abundance of PGA^+^ and EPS^+^ CFUs from 0% to 100% (see Methods 3). This approach allowed us to traverse the morphological biofilm phase space defined by the joint contributions of PGA and EPS. We characterized colony morphology across the phase space using both side-view OCT cross-sections (Fig. 5c) and top-down stereomicroscopy images (Fig. 5d). Biofilms with low levels of either PGA or EPS formed smooth colonies, while the addition of either of the other ECM polymer was sufficient to induce wrinkling. Biofilms in regions with high CFU fractions of both polymers exhibited pronounced wrinkling.

To quantify the morphologies we observed in our co-cultures, we measured thickness and wrinkling. We determined thickness across the morphological phase space by measuring the maximum biofilm thickness from segmented OCT cross-sections (Fig. S3a). We used maximum rather than mean thickness because swollen, highly wrinkled colonies also contained very thin regions. The result is plotted as a heatmap in Fig. 5e. Maximum thickness increased primarily with the fraction of PGA producers, while EPS had minimal influence on vertical growth. This observation is consistent with PGA’s role in driving water uptake and swelling. To quantify the degree of surface wrinkling, we introduced a dimensionless coefficient, *q*, defined as the fraction of the biofilm area occupied by wrinkles (*q* = Wrinkle area*/*Biofilm area; see Methods; Fig. S3b). This quantity would be zero for highly smooth biofilms. The heatmap of *q* (Fig. 5f) captures a distinct morphological transition as matrix composition changes. When either PGA or EPS production is low, biofilms remain smooth (*q* ≈ 0). However, as both polymer fraction is increased, the surface becomes increasingly wrinkled, giving rise to a sharp transition in *q* with a phase-space boundary consistent with that of our model’s phase diagram (Fig. 5b). This experimental behavior supports our hypothesis that a cooperative interaction between PGA and EPS is necessary for wrinkling. While PGA drives swelling-induced in-plane compression, EPS cross-links biomass to provide biofilm elasticity that supports mechanical stresses and leads to buckling and wrinkle formation. Varying the production of the two polymers explores a diverse phase space of biofilm morphologies.

## Discussion

In this study, we identify two self-secreted polymers, poly-*γ*-glutamic acid (PGA) and exopolysaccharides (EPS), whose synergistic effect on biofilm material properties drives *B. subtilis* colony morphogenesis. PGA induces water absorption and EPS effectively cross-links biomass, causing gelation. Together, PGA and EPS turn biofilms into swelling gels that develop large wrinkles. By modulating polymer production with co-cultures of matrix knockouts, we move across a morphological phase space between smooth and wrinkled colonies as predicted by a simple bilayer wrinkling model. Our results establish a mechanistic link between molecular-scale polymer production and colony-scale morphology and show that two parameters can predictably modulate the highly complex process of biofilm morphogenesis.

Our work uses *B. subtilis* as a powerful and controllable system with which to explore how active and living systems harness soft matter physics to achieve desired functionality [42–44]. Specifically, we have revealed a self-driven phase transition for which a passive elastic model can provide insight but is unable to capture the highly dynamic changes over the colony life cycle. We hope that our findings will inspire new work to investigate the origins and consequences of active, self-driven phase transitions in cellular communities.

Our results raise many questions about the physics of biofilm morphogenesis in PGA-producing strains, which appears to be fundamentally different from wrinkle formation in low-PGA strains, such as the typical 3610[6, 45]. Our model rationalizes wrinkling vs. non-wrinkling based on mechanical strain and stiffness, but we anticipate that biofilm morphology depends on other factors, such as cell growth, surface adhesion, and water absorption, all of which vary with the timing of PGA and EPS production. More detailed modeling and biophysical experiments will reveal how biofilm chemomechanics across the PGA-EPS phase space regulate colony topography.

We focus on *B. subtilis* strain 3610 pBS32^0^ because PGA production has a dramatic impact on its biofilm physics. The vast majority of *B. subtilis* biofilm research has examined 3610, where PGA production has no impact on biofilm morphology[46]. However, this is likely due to 3610’s *rapP* quorum sensing mutation that inhibits PGA production[25]. Even in 3610, however, Morris, et al. showed that under conditions where the strain does produce PGA, biofilm expansion slows during PGA production and is restored by EPS production[15], consistent with EPS gelating a PGA-rich liquid-like environment during biofilm growth. Numerous strains that lack 3610’s mutated *rapP* gene exhibit the same thick, wrinkled morphology we observe in our WT[47]. We believe that this ubiquitous morphology arises from the swelling gel formation we explore in this paper. Given that this biofilm morphology is strongly conserved, it is important to ask what the function of gel formation may be in the soil environment of *Bacillus*. High PGA production is also directly coupled to high rates of biofilm spore formation[25]. Gel formation may consequently play a role in spore dispersal or germination, for example by osmotically absorbing nutrients and/or releasing spores during moisture spikes in the soil. Furthermore, forming PGA-rich gels could allow biofilms to retain water for survival during dry periods[48].

In our co-culture experiments, we observed relatively stable genotypic proportions. Typically in such experiments, genotypes spatially segregate into sectors during colony growth[49, 50]. Our co-culture biofilms, however, appear to be homogeneous materials. Wrinkles are not patchy and height profiles are generally smooth (5d). These observations suggest that EPS and PGA are approximately evenly distributed throughout the biofilm, either because the EPS and PGA diffuse to uniformity before becoming cross-linked or because the EPS- and PGA-producing cells are well-mixed. It is possible that fluid formation near the growing biofilm exterior facilitates mixing that inhibits genotypic fixation and sectoring. It will be valuable to investigate spatial distributions of genotypes to determine the effect of gelation on community ecology.

## Methods

### Bacterial strains

The primary strain of *B. subtilis* used in this study is the common biofilm strain NCIB3610, cured of the pBS32 plasmid. This plasmid-free strain pBS32^0^, is more transformable due to its lack of the plasmid-borne gene *comI*, which inhibits genetic competence [24]. pBS32 also encodes RapP, a quorum sensing gene whose pBS32 allele inhibits both PGA production and spore formation[25]. The three other strains used are the three gene knockouts in 3610 pBS32^0^: Δ*eps* which produces PGA but not EPS, Δ*pgsB* which produces EPS but not PGA, and the double knockout Δ*pgsB* Δ*eps*, which makes neither of the polymers (Table 1).

**Table 1.** Base strains used in this study. All strains are derived from *B. subtilis* NCIB3610 pBS32^0^.

| Strain | Matrix phenotype | Genotype |
| --- | --- | --- |
| WT | PGA<sup>+</sup>, EPS<sup>+</sup> | NCIB3610 pBS32<sup>0</sup> plasmid-free (3610PF) |
| $\Delta pgsB$ | PGA<sup>-</sup>, EPS<sup>+</sup> | 3610PF *pgsB::cat* |
| $\Delta eps$ | PGA<sup>+</sup>, EPS<sup>-</sup> | 3610PF *epsA-O::tet* |
| $\Delta pgsB \Delta eps$ | PGA<sup>-</sup>, EPS<sup>-</sup> | 3610PF *pgsB::cat epsA-O::tet* |

### Growth conditions and biofilm formation

All biofilms were grown using the same protocol. The strains were streaked on an LB plate and kept overnight in a 37^◦^C incubator. One small, circular colony was isolated and grown in 1mL liquid LB up to an OD_600_ of ∼0.6. At this point, the culture was centrifuged for 3 minutes at 5000 rpm and cells were resuspended in MSgg solution [5 mM potassium phosphate (pH 7.0), 100 mM 3-morpholinopropane-1-sulfonic acid (pH 7.0), 0.5% glycerol, 0.5% glutamate, 700 *µ*M CaCl_2_, 2 mM MgCl_2_, 50 *µ*M MnCl_2_, 100 *µ*M FeCl_3_.6H_2_O, 1 *µ*M ZnCl_2_, 2 *µ*M thiamine HCl]. The MSgg was culture was grown in a 37^◦^C shaker for an additional ∼1 hour to an OD_600_ of ∼0.5 (cultures that grew to a higher value were diluted to OD_600_ of ∼0.5 with MSgg solution). 1*µ*L of the MSgg culture was spotted on a MSgg-supplemented 2% w/v agarose pad and incubated upside down at 30^◦^C for the entire duration of the experiment.

### Stereoscope imaging

Stereoscope images of bacterial biofilms were taken using a 1X objective of an Olympus SZX7 stereomicroscope equipped with an overhead ring lamp (Fig. 5e and Fig. 1a-d). The glossy appearance in the high fluid content biofilms can be attributed to the reflection from the ring light in high water-containing biofilms. Glare from the illumination light has been removed from Fig. 1c by segmenting out the reflection and filling in the area with cv2.inpaint.

### OCT imaging and analysis

2D *xz* cross-sectional scans of dimension 5 × 1 mm and 3 *µ*m/pixel resolution were obtained with a Thorlabs Telesto OCT system. The surfaces of the biofilm were segmented using the following steps: contrast stretching to normalize the intensity histogram, Otsu thresholding to extract the bright features, binary opening to remove small objects and an additional small object removal step, and finally defining the highest non-zero pixel along *z*-direction as the maximum biofilm height. To obtain the baseline corrected surface profiles, a 2-degree (for flat biofilms that stick to the agarose) or 1-degree (for wrinkled biofilms where baseline correction is needed to primarily fix rotations) polymeric fit was subtracted from the raw surface data. The corrected surface profiles from *n* = 3 replicates were used to obtain the averaged maximum height heatmap in Fig. 5e. One of the three replicates with corrected surfaces is shown in Fig. S3a.

### Time-lapse phase imaging and cellular motion tracking

Phase-contrast imaging was done on an Olympus IX83 microscope with the 10X, 0.3NA objective. For particle trajectory analysis, 60-second videos were captured at 0.87 frames/s at for biofilms after 0 and 20 hours of growth for the four base strains (Table 1) (Videos V1-V4) and also for the WT strain at *t* = 0 hours before and after the initial spot dried as positive and negative control (Videos V5, V6). The imageJ TrackMate plug-in[51] was used to track the cellular features in the phase videos. A Difference of Gaussian (DoG) detector with an estimated object diameter of 1*µ*m was set to detect cell-sized features. Nearest neighbor tracking with a maximum frame-to-frame linking distance set to 10*µ*m yielded the tracked trajectories, which have been color-coded based on the mean speed of the tracked cells. In Fig. 2e-h and Fig. S2b, c, e-h, the pink spots are the initial locations of the cells tracked.

### Co-culture construction

To construct biofilms having specific fractions of PGA and EPS producers, we made co-cultures of the four base strains that span all the possible combinations of polymer production (Table 1). To make a co-culture, the two strains were grown following the procedure described above, each grown/diluted to an OD_600_ value of 0.5 (to ensure one-to-one mapping between volume and number of cells), and then mixed in the desired ratio. For example, to make a biofilm in which 50% of the cells make PGA and EPS the other 50% does not make any of the polymers, we mixed strains *WT* and Δ*pgsB* Δ*eps* in equal parts, and then spotted 1*µL* of the mixed co-culture on a 1X Msgg-supplemented 2% agarose pads and incubated upside down (to prevent condensation droplets) at 30^◦^C for the duration of the experiment. The phase portrait of co-cultures is constructed by mixing the four base strains in volume ratios shown in Fig. S5.

### Antibiotic CFU plating assay

To assess whether the relative abundances of different strains were maintained at their initial co-culture mixing ratios, we grew co-culture biofilms following the protocol described in the Methods section . Co-cultures were plated for colony-forming unit (CFU) counts using a serial dilution assay on LB agar plates supplemented with the appropriate antibiotics (Table 2). For day 0 CFU counts, 25*µL* of the initial co-culture mixture at *OD*_600_ ≈ 0.5 was used. For day 1 and day 2 counts, biofilms were carefully scraped from the surface of MSgg-supplemented agarose pads and resuspended in 400*µL* of 1 × PBS. Plates were incubated overnight at 30^◦^*C*, and colonies were counted to determine total CFUs. Relative abundance was calculated as the ratio of one strain to the other, across three time points (day 0–2) and three initial mixing ratios and three replicates, as shown in Fig. 3c and 4f.

**Table 2.** Antibiotic CFU plating assay.

| Co-culture | Antibiotic plates used |
| --- | --- |
| $\Delta eps:\Delta pgsB \Delta eps$ | *tet* - $\Delta eps$ and $\Delta pgsB \Delta eps$ , *cat*-$\Delta pgsB \Delta eps$ |
| $\Delta pgsB:\Delta pgsB \Delta eps$ | *cat* - $\Delta pgsB$ and $\Delta pgsB \Delta eps$ , *tet*-$\Delta pgsB \Delta eps$ |
| WT:$\Delta pgsB$ | *spec* - WT, *cat*-$\Delta pgsB$ |
| WT:$\Delta eps$ | *spec* - WT, *tet*-$\Delta eps$ |

### Water dissolution assay

To test the resistance of the biofilm to water addition and to obtain the fraction of biofilm resistant to dissolution, the biofilms were submerged in water by pipetting a fixed volume (∼3 mL) of deionized water into the wells containing the biofilms. The system was manually perturbed by a slight in-plane shaking for 1 minute and then allowed to rest for 30 minutes (Fig. 4d). The area of the biofilm before and after the water dissolution is obtained by manual thresholding of biomass from the images taken on a Nikon 7500 digital camera, and the corresponding retention area ratio is computed (Fig. 4c, e). For obtaining the fold change in height, we used a DataPhysics OCA 25 contact angle goniometer (Fig. 4c).

### Wrinkling coefficient analysis

The extent of wrinkling is quantified by defining a dimensionless wrinkling coefficient *q*, which is the fraction of the biofilm area that is wrinkled:

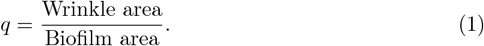

The wrinkle area and biofilm area for 25 biofilms in the phase portrait are obtained by manually thresholding grayscale stereoscope images (Fig. 5c) to obtain the wrinkles and the biomass area (in pixel^2^) in ImageJ (Fig. S3b). The analysis is averaged over three replicates to obtain the wrinkling coefficient heatmap shown in Fig. 5e.

## Acknowledgements

We acknowledge funding from NIH award No. R35GM142584 (J.W.L.) and a Burroughs Wellcome Fund Career Award at the Scientific Interface (J.W.L.). This research was supported by the Biomedical Engineering Core Facilities at Boston University. The authors acknowledge the use of the facilities of the Boston University Neurophotonics Center.

## Declarations

The authors declare no competing financial interests.

## Code Availability

Code is available at https://github.com/Larkin-Lab/saha_et_al_gel_biofilm

## Supplementary Information

### Supplementary Videos

- Video V1: Δ*pgsB* Δ*eps* 10X phase time-lapse video at t = 10 hours.
- Video V2: Δ*pgsB* 10X phase time-lapse video at t = 10 hours.
- Video V3: Δ*eps* 10X phase time-lapse video at t = 10 hours.
- Video V4: WT 10X phase time-lapse video at t = 10 hours.
- Video V5: Wet droplet 20X phase time-lapse video.
- Video V6: Dry droplet 20X phase time-lapse video.

### Thin-film bilayer model

We modeled biofilms as thin films of Young’s modulus *E*_*f*_ adhered to agarose substrates of Young’s modulus *E*_*s*_ [7, 36]. For such a bilayer system under strain, film wrinkling will occur beyond a critical strain measured relative to the stress-free configuration *ϵ*_*c*_ ∼ (*E*_*s*_*/E*_*f*_) ^2*/*3^ [37–39, 52]. Because all the biofilms were grown on 2% agarose substrates, we assume that *E*_*s*_ is a constant. Thus, the critical strain necessary for the emergence of wrinkles in this simplified model is solely modulated by the biofilm stiffness *E*_*f*_ . For a biofilm to wrinkle, the strain, *ϵ*, must exceed *ϵ*_*c*_. PGA or EPS could cause wrinkling by either increasing *ϵ* or decreasing *ϵ*_*c*_ (via *E*_*f*_). In our experiments, PGA causes water absorption and swelling without a significant impact on the extent of dissolution in water (Fig. 3, Fig. 4), while EPS prevents dissolution without causing significant swelling (Fig. 2, Fig. 4). For these reasons, we assume that strain depends primarily on the fraction of PGA-producers, *f*_*PGA*_, and that stiffness depends primarily on the fraction of EPS-producers, *f*_*EPS*_. In the following sections, we discuss how PGA-induced swelling modulates in-plane strain and EPS-driven gelation modulates the critical strain necessary for the emergence of wrinkles.

### PGA-induced isotropic swelling gives rise to in-plane strain

Through our experiments, we demonstrate that PGA production promotes fluid uptake from the underlying substrate, resulting in a monotonic increase in the biofilm’s maximum height with the fraction of PGA-producing cells. Concurrently, we observe a reduction in the base radius as the concentration of PGA polymers increases (Fig. S7a-c). To interpret how these morphological changes generate in-plane strain, we propose a simple physical model: envision water being injected into a sponge whose base is constricted and moderately adhered to the substrate, causing isotropic swelling. In this scenario, an increase in water volume, together with the fixed base, produces an in-plane compressive strain. Assuming that the volume of absorbed fluid – and hence the strain – is a monotonically increasing function of the fraction of PGA polymers (*f*_*PGA*_), we can express the strain with the following scaling relation:

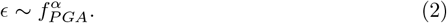

We require the power law exponent *α >* 0 so that *ϵ* increases with increasing *f*_*PGA*_, but leave its value unspecified to maintain generality. The key assumption made here is that the EPS polymer does not contribute to the in-plane strain. Based on our data, this appears to be a reasonable assumption because neither the height nor the biofilm footprint, indirect indicators of in-plane strain, are found to change significantly when the EPS fraction is varied in experiments at a fixed PGA fraction.

### EPS drives sol-gel phase transition via cross-linking

In our system, EPS-producing cells generate cross-linking EPS polymer that drives a gelation phase transition from a sol state at low EPS fractions to a gel state at high concentrations of EPS polymers. As shown in Fig. 4e, addition of water to the biofilm reveals a phase transition in which the fraction of retained biofilm area, a proxy for gel fraction *P*_*gel*_ i.e., fraction of connected EPS clusters, increases sharply and follows a power law:

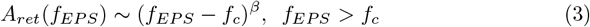

with experimentally determined values *β* = 0.271 ± 0.114 and *f*_*c*_ = 0.126 ± 0.078 by fitting the area retained on water addition as a function of *f*_*EPS*_ plot to the proposed power law (Fig. S7d). Below the threshold, the biofilm fails to resist dissolution, indicating the absence of a coherent gel network.

Gelation is often modeled as a percolation transition [32]. A key result from these theories is that the emergence of mechanical rigidity in polymer networks is intimately linked to the formation of a system-spanning cluster of cross-links. Theoretical and experimental studies of polymer gels and percolation networks have established that the onset of elasticity coincides with the formation of the percolating cluster that marks the gel point [53, 54]. In these models, the elastic modulus emerges only above the critical fraction *f*_*c*_ where long polymer chains get strongly cross-linked, forming an elastic gel material. Below the threshold *f*_*c*_, the material behaves like a liquid, having little to no elasticity. Hence, we expect the Young’s modulus *E*_*f*_ to follow a scaling relation:

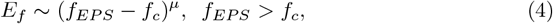

where the exponent *µ* characterizes the steepness of the modulus growth near the threshold and is typically expected to be larger than *β* [55, 56]. The power-law scaling of Young’s modulus above the EPS fraction threshold *f*_*c*_ is a well-established feature of percolation and gelation transitions, and is consistent with our measurements of retained biofilm area. In three dimensions, the elasticity scaling exponent *µ* takes values of 3 in the mean-field model, 1.8 in the electrical analogy model [57], and 3.7 in the bond bending model [58]. Experimentally, *µ* has been observed to extensively vary, with values near 1.7 for gelatin [59], ∼1.9 measured in diisocianate-triol gels [60] and approaching 3 in polyesters [61]. Accordingly, we consider a wide range of critical exponents *µ* ≈ 1.5-4, to test the robustness of our model. The key assumption made here is that the stiffness modulus *E*_*f*_ of the biofilm scales independently of the PGA fraction *f*_*PGA*_. This is consistent with our observation that the Δ*eps* strain, which is devoid of any EPS polymer and solely has PGA polymer producers(PGA^+^, EPS^−^), dissolves in water and behaves like a sol because it lies below the critical crosslinking threshold *f*_*c*_ necessary for the sol-gel transition.

### Phase boundary between wrinkled and smooth surface morphology

Combining these relationships between the PGA swelling-induced strain (eq. 2), the EPS-driven gelation phase transition and stiffening (eq. 4) and exploiting the equation for critical strain necessary for wrinkling in the elastic bilayer model 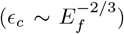, we can predict the boundary where biofilms transform from smooth to wrinkled, i.e., where the strain *ϵ* generated by swelling from the PGA-producers at fraction *f*_*PGA*_ equals the critical strain *ϵ*_*c*_, which in turn is determined by the fraction *f*_*EPS*_ of EPS polymers. At this boundary, we rewrite the expression for strain in terms of the *f*_*PGA*_ and that for Young’s modulus in terms of *f*_*EPS*_ and the other model parameters *f*_*c*_, *α*, and *µ* to obtain the equation for the phase boundary as follows:

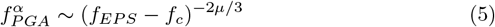

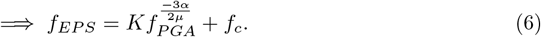

Here *K* is a proportionality constant. Fig. S8a, b shows an agreement between the experimental wrinkling phase portrait and the one derived from the model by setting *α* = 2, *µ* = 1.5 and *K* = 0.01. Fig. S8c shows a qualitative agreement of our experimental result with this theoretical model over a wide range of model parameters *α* and *µ*.

---

**Fig. S1.**
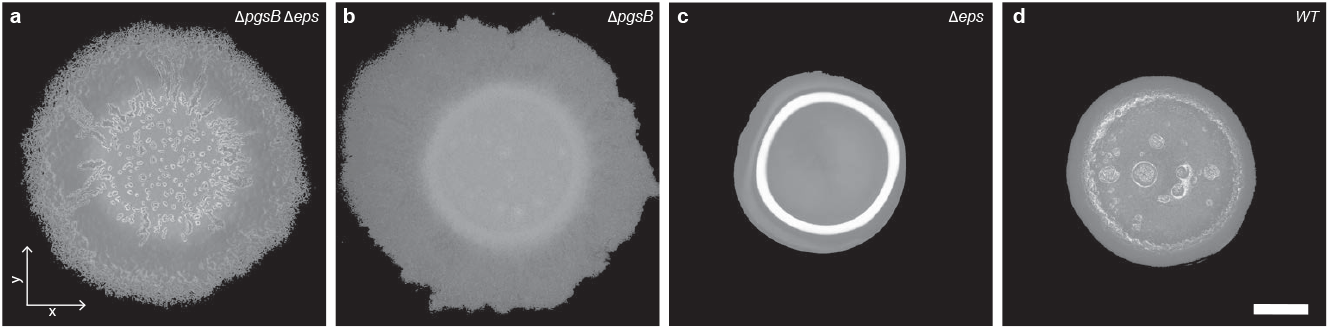
Top-down stereoscopic images of 24-hour-old biofilms formed by strains with differential production of PGA and EPS polymers. **a**, Δ*pgsB* Δ*eps* (neither polymer), **b**, Δ*pgsB* (EPS only), **c**, Δ*eps* (PGA only), **d**, WT (both polymers). The bright ring in **c** is a reflection of the overhead illumination light due to the excessively shiny surface of Δ*eps* colonies.

**Fig. S2.**
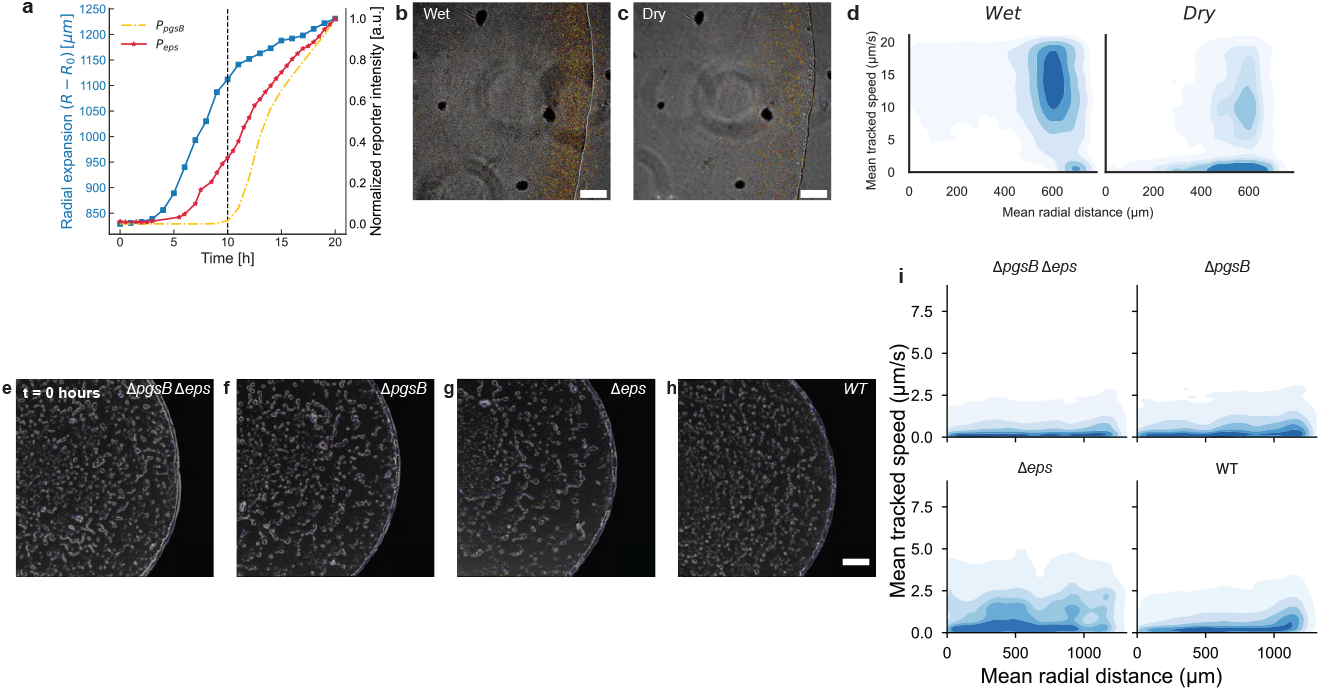
Reporter dynamics and spatial distribution of cell speeds in biofilms (a, e–i) and in a liquid droplet (b–d) during the wet–dry transition. **a**, Colony radius (blue) of the wild-type (WT) strain over 20 hours of growth, together with constitutive reporter activity (*P*_*pen*_, red) and the reporter for the PGA biosynthesis gene *pgsB* (*P*_*pgsB*_, yellow). Signal from the *pgsB* reporter emerges at *t* ≈ 10 hours. **b,c**, Phase-contrast images (20 ×) of WT droplets immediately after spotting in the wet state (**b**) and following drying (**c**). Scale bar, 100 *µ*m. **d**, Kernel density estimates (KDEs) of mean tracked cell speeds and radial trajectories in wet versus dry droplets reveals a shift from nonzero velocities in the wet state to near-zero speeds upon drying. **e-h**, Phase-contrast images (10 ×) of Δ*pgsB* Δ*eps*, Δ*pgsB*, Δ*eps* and WT at *t* ≈ 0 hour. Scale bar, 200 *µ*m. **i**, Kernel density ≈ estimates (KDEs) of mean tracked cell speeds and radial trajectory distributions at *t* ≈ 0 hour show uniformly scattered, low-speed motion across all strains.

**Fig. S3.**
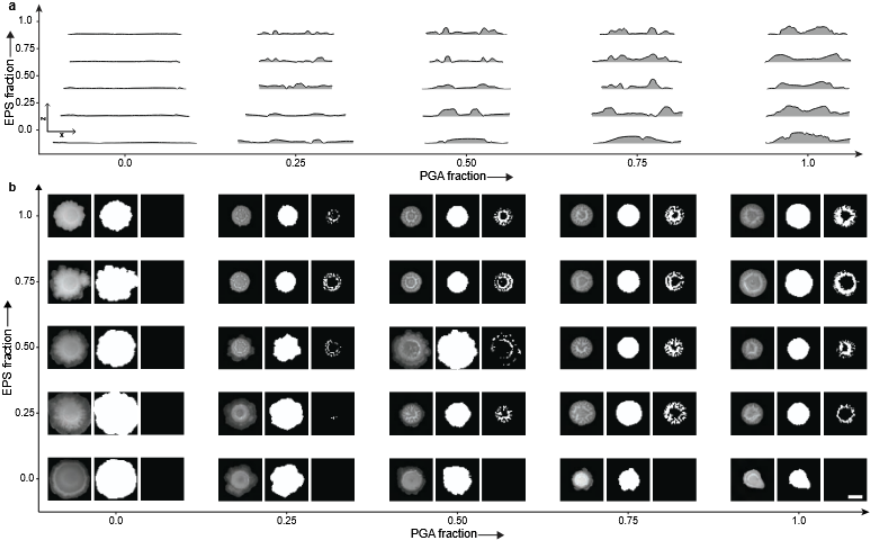
Phase diagrams of segmented OCT *xz* profiles and wrinkle/biomass masks. **a**, Phase diagram of segmented surfaces from optical coherence tomography (OCT) *xz*-profiles of biofilms with increasing fractions of PGA and EPS–producing cells (0–100%). Scale bar 1 mm. **b**, Phase diagram showing wrinkling coefficient analysis workflow - stereoscopic grayscale image and the corresponding biomass and wrinkle masks(left to right).

**Fig. S4.**
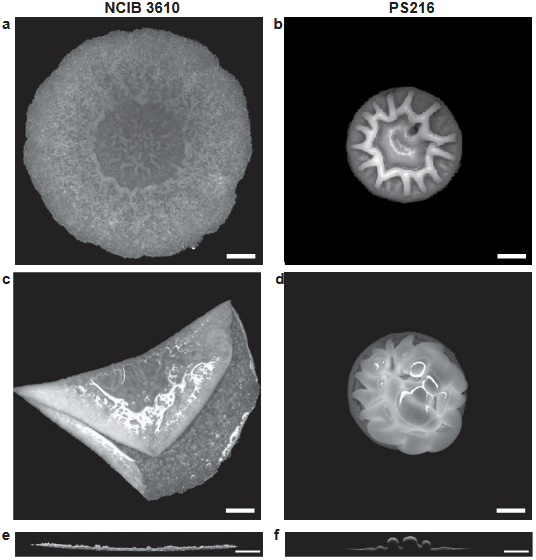
Top-down stereoscopic images and OCT *xz* profiles of NCIB3610 and PS216 biofilms before and after water immersion. **a-d**, Top-down stereoscopic images of 48-hour-old *Bacillus subtilis* biofilms formed by strains NCIB3610 and PS-216 before (**a, b**) and after water immersion (**c,d**). Scale bar, 2 *mm*. NCIB3610 does not produce PGA under standard biofilm conditions due to a mutation in the plasmid-borne gene *rapP*. PS216 naturally lacks the pBS32 plasmid that carries *rapP*. We expect it to behave similarly to our WT, 3610 pBS32^0^. **e, f** Corresponding *xz* profiles measured by optical coherence tomography. Scale bar, 1 *mm*.

**Fig. S5.**
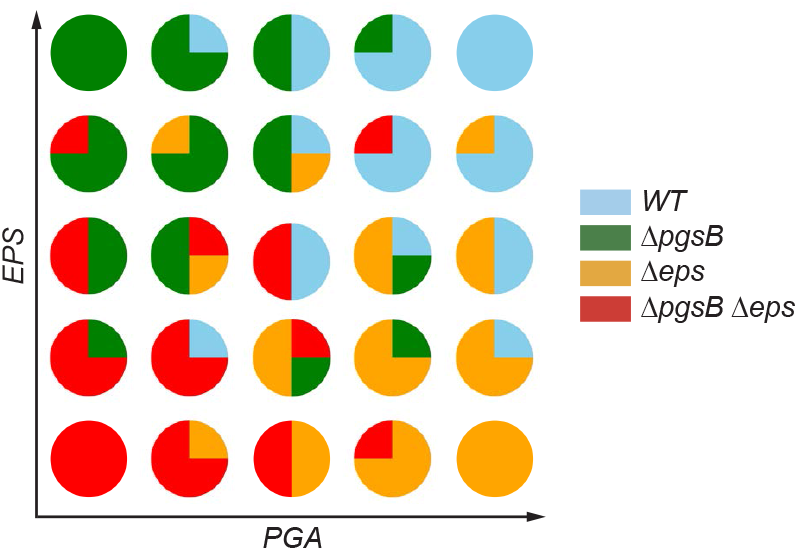
Pairwise combinations of four Bacillus subtilis strains visualized as pie charts representing their relative contributions in co-culture and co-co-culture assays. Each pie chart depicts the proportion of two/three strains in the mixture, with colors corresponding to strain identities: WT (light blue), Δ*pgsB* (green), Δ*eps* (gold), and Δ*pgsB* Δ*eps*(red). The *x*-axis (PGA) and *y*-axis (EPS) indicate increasing producer fractions of produce poly-*γ*-glutamate or exopolysaccharide, respectively.

**Fig. S6.**
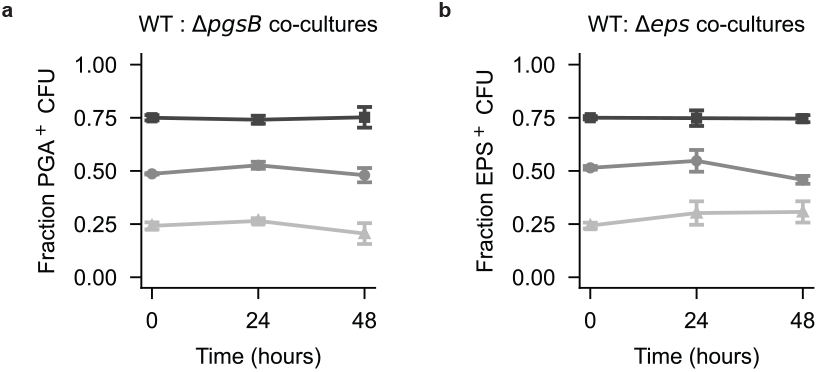
Proportion of PGA-producing and EPS-producing CFUs over time in co-culture biofilms. **a, b** Temporal evolution of PGA^+^ CFU fraction in a co-culture of WT : Δ*pgsB* (**a**) and of EPS^+^ CFU fraction in a co-culture of WT : Δ*eps* (**b**) with three different initial genotypic ratios of Δ*pgsB* and Δ*pgsB* Δ*eps* cells, 0.75:0.25 (squares), 0.5:0.5 (circles), and 0.25:0.75 (triangles). Points represent average ± S.D. from three independent replicates (*n* = 3).

**Fig. S7.**
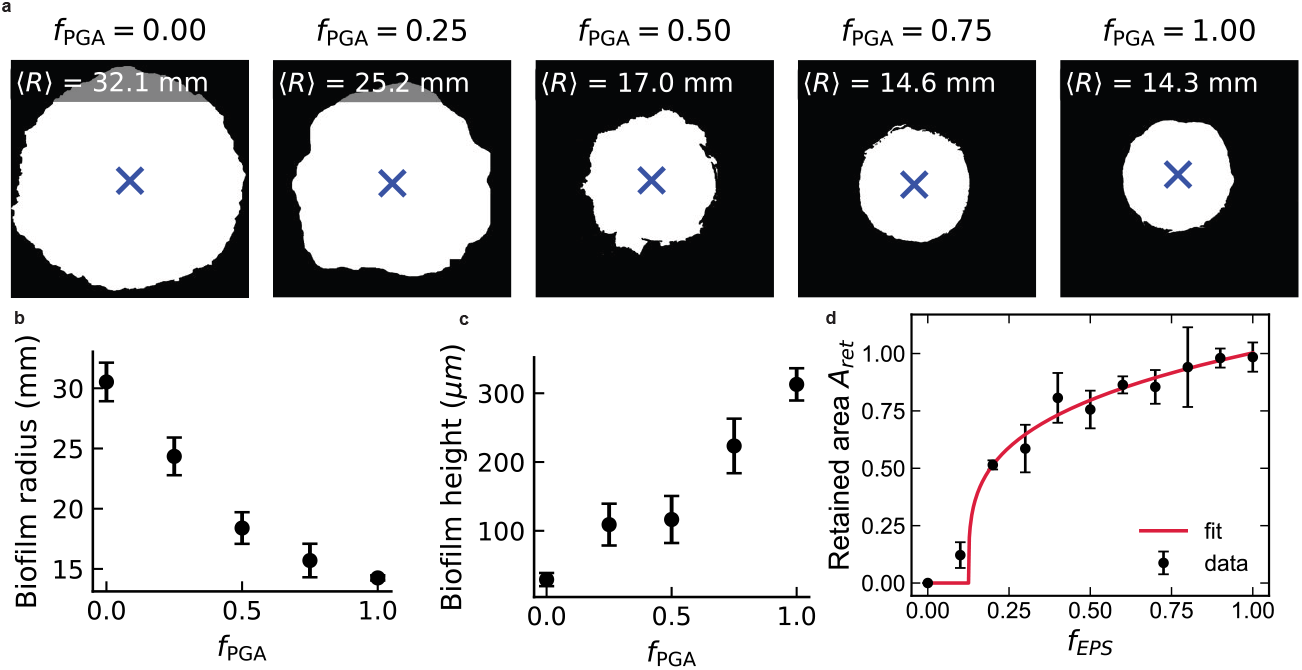
Representative binary masks, colony metrics versus PGA fraction and retained area versus EPS fraction. **a**, Representative binary masks of biofilm colonies at increasing PGA fractions (*f*_PGA_), showing colony centroids (blue crosses) and mean colony radii (*R*). **b**, The mean biofilm radius *R* decreases as *f*_PGA_ increases. **c**, The maximum biofilm height *h* increases monotonically with *f*_PGA_. **d**, Retained area *A*_ret_ as a function of EPS fraction *f*_*EPS*_, showing experimental data (black circles, mean ± S.D. from three independent replicates (*n* = 3)) and fit to the theoretical model (*f*_*EPS*_ − *f*_*c*_)^*β*^ with *β* = 0.271 ± 0.114 and *f*_*c*_ = 0.126 ± 0.078 (red line).

**Fig. S8.**
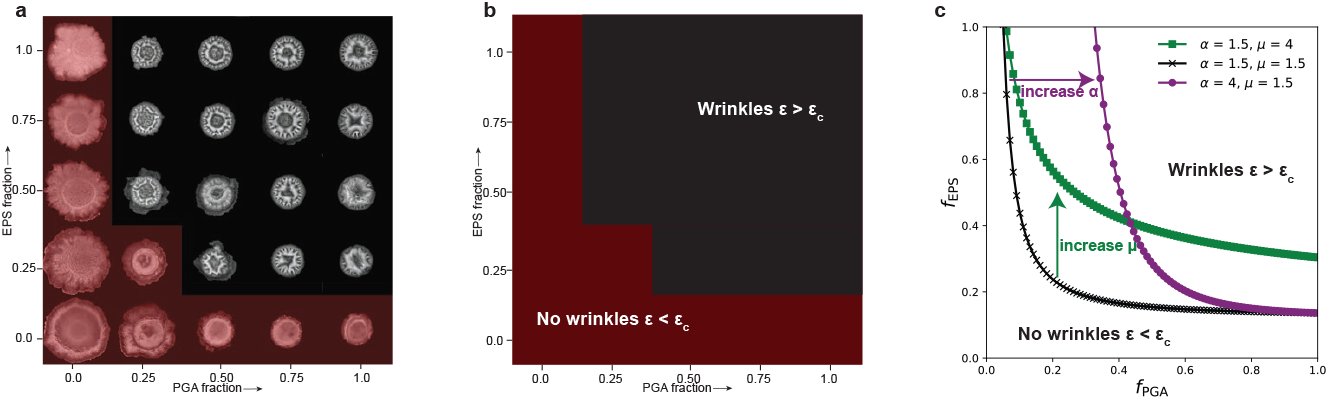
Comparision between experimental and model phase portraits. **a**, Phase portrait of biofilm morphologies across PGA and EPS fractions, showing representative images for each condition, overlaid with a red layer for the smooth biofilms. **b**, Binary phase portrait generated from the model with *α* = 2, *µ* = 1.5, *f*_*c*_ = 0.126 and *K* = 0.01. **c**, Phase boundaries obtained from the model for a range of parameters *α* and *µ* ∈ [1.5, 4]

## Notes

https://github.com/Larkin-Lab/saha_et_al_gel_biofilm

### Competing Interest Statement

The authors have declared no competing interest.

### Summary of Updates

Errors corrected in the references and language clarified in the modeling section.

